# Methionine oxidation of Carbohydrate-Active enZymes during white-rot wood decay

**DOI:** 10.1101/2023.10.24.563746

**Authors:** Lise Molinelli, Elodie Drula, Jean-Charles Gaillard, David Navarro, Jean Armengaud, Jean-Guy Berrin, Thierry Tron, Lionel Tarrago

## Abstract

White-rot fungi employ secreted carbohydrate-active enzymes (CAZymes) along with reactive oxygen species (ROS), like hydrogen peroxide (H_2_O_2_), to degrade lignocellulose in wood. H_2_O_2_ serves as a co-substrate for key oxidoreductases during the initial decay phase. While the degradation of lignocellulose by CAZymes is well-documented, the impact of ROS on the oxidation of the secreted proteins remains unclear and the identity of the oxidized proteins is unknown. Methionine (Met) can be oxidized to Met sulfoxide (MetO) or Met sulfone (MetO_2_) with potential deleterious, antioxidant, or regulatory effects. Other residues, like proline (Pro), can undergo carbonylation. Using the white-rot *Pycnoporus cinnabarinus* grown on aspen wood, we analyzed the Met content of the secreted proteins and their susceptibility to oxidation combining H ^18^O with deep shotgun proteomics. Strikingly, their overall Met content was significantly lower (1.4%) compared to intracellular proteins (2.1%), a feature conserved in fungi but not in metazoans or plants. We evidenced that a catalase, widespread in white-rot fungi, protects the secreted proteins from oxidation. Our redox proteomics approach allowed identification of 49 oxidizable Met and 40 oxidizable Pro residues within few secreted proteins, mostly CAZymes. Interestingly, many of them had several oxidized residues localized in hotspots. Some Met, including those in GH7 cellobiohydrolases, were oxidized up to 47%, with a substantial percentage of sulfone (13%). These Met are conserved in fungal homologs, suggesting important functional roles. Our findings reveal that white-rot fungi safeguard their secreted proteins by minimizing their Met content and by scavenging ROS and pinpoint redox-active residues in CAZymes.

**Importance:** The study of lignocellulose degradation by fungi is critical for understanding the ecological and industrial implications of wood decay. While carbohydrate-active enzymes (CAZymes) play a well-established role in lignocellulose degradation, the impact of hydrogen peroxide (H_2_O_2_) on secreted proteins remains unclear. This study aims at evaluating the effect of H_2_O_2_ on secreted proteins, focusing on the oxidation of methionine (Met). Using the model white-rot model *Pycnoporus cinnabarinus* grown on aspen wood, we showed that fungi protect their secreted proteins from oxidation by reducing their Met content and utilizing a secreted catalase to scavenge exogenous H_2_O_2_. The research identified key oxidizable Met within secreted CAZymes. Importantly, some Met like those of GH7 cellobiohydrolases, undergone substantial oxidation levels suggesting important roles in lignocellulose degradation. These findings highlight the adaptive mechanisms employed by white-rot fungi to safeguard their secreted proteins during wood decay and emphasize the importance of these processes in lignocellulose breakdown.

## Introduction

Filamentous fungi are crucial in the carbon cycle because of their ability to decompose organic matter and release carbon back into the ecosystem (1). Wood-decay fungi extract carbon from recalcitrant polymers like cellulose and hemicelluloses embedded in a lignin matrix, by secreting enzymes into the extracellular environment (2). The secreted proteins, commonly referred to as the “secretome”, contains the carbohydrate-active enzymes (CAZymes), including large sets of glycoside hydrolases (GH) and auxiliary activity (AA) enzymes (3, 4). The GH class contains the endo– and exo-acting enzymes targeting cellulose and hemicelluloses. The AA class contains laccases (AA1) and class II peroxidases (AA2) involved in lignin degradation, and lytic polysaccharide monooxygenases (LPMO, AA9) proposed to play an important role in the early phase of wood degradation (5–7). Reactive oxygen species (ROS), particularly hydrogen peroxide (H_2_O_2_), play a crucial role in the early phase of wood decay (8–10). H_2_O_2_ is generated by glucose-methanol-choline oxidoreductases (AA3) and copper radical oxidases (AA5), that target carbohydrate or aliphatic/aromatic alcohols (11, 12). In white-rot fungi, H_2_O_2_ acts as a co-substrate to fuel different types of LPMOs and AA2 peroxidases (13–15). While the roles of CAZymes and H_2_O_2_ during the degradation of lignocellulose is well-documented, the impact of ROS on the oxidation of CAZymes remains unclear. Fenton-generated ROS were shown to decrease enzymatic activity of some GHs in white-rot secretomes, through the oxidation of the secreted proteins, and the most abundant form of oxidation found was the oxidation of Met residues (16). Met can be converted by H_2_O_2_ into Met sulfoxide (MetO) or Met sulfone (MetO_2_) and this has been shown to play critical roles in many organisms (17–20). For instance, Met oxidation can lead either to a loss of function or act as a regulatory switch, to activate enzymes (21, 22). The presence of oxidizable Met in proteins can also be associated to an antioxidant role by serving as a protection to avoid the oxidation of residues critical for the protein function (23). MetO can be reduced back to Met by the methionine sulfoxide reductases, but, as these enzymes are most likely not secreted, they cannot reduce oxidized Met of secreted proteins (24). Regarding the importance of H_2_O_2_ during wood decay, our hypothesis is that some Met of secreted proteins can be oxidized and play antioxidant or regulatory roles. Interestingly, the treatment of an α-galactosidase (GH27) from *Trichoderma reesei* with H_2_O_2_ strongly increased its activity due to the oxidation of a Met located near the active site, suggesting that Met oxidation of CAZymes can have a positive effect on their activity (25). To identify secreted proteins with oxidizable Met, we selected the white-rot *Pycnoporus cinnabarinus* grown on aspen wood as a model. *P. cinnabarinus* secretes H_2_O_2_-producing enzymes in the early phase of wood degradation, to provide co-substrate for the H_2_O_2_-consuming CAZymes (26), but also as potential oxidant of Met. We characterized the secreted proteins and identified their oxidized Met with a redox proteomics approach using ^18^O-labeled hydrogen peroxide (H ^18^O) as a blocking reagent to prevent artifactual oxidation (27–30). Our investigations reveal the extend of Met oxidation in fungal secretomes and the precise localization of oxidizable residues in secreted proteins and CAZymes, suggesting potential antioxidant or regulatory roles.

## Results

### Secreted proteins retrieved by ion exchange chromatography are representative of early days decay

To identify oxidized Met in the secreted proteins during *P. cinnabarinus* growth, we focused on the early days of wood decay, during which production and consumption of H_2_O_2_ are likely to occur (26). We performed a time-course growth of the fungus on aspen wood and purified the proteins from the culture supernatants at days 3, 5 and 7. To avoid artifactual oxidation that can occur with standard methods combining a freezing step to protein concentration with ultrafiltration (26, 31, 32), we purified the secreted proteins on successive anion and cation exchange chromatography. Then, we oxidized the secreted proteins with H_2_^18^O_0_ and identified them and their potential oxidative Met modifications using proteomics (**Fig. S1**). We identified 308 secreted proteins, among which CAZymes and proteases represented 41% and 10%, respectively (**Fig. 1**; **Table 1; Dataset 1**). Among the CAZymes, GH was by far the most represented class, with 89 proteins (71%) spread over 34 families (**Fig. 1; Table 1**). We identified most of the CAZymes expected during wood degradation, i.e. cellulases (GH3, GH5_5, GH5_9 and GH7, GH12, GH45), hemicellulases (GH10, GH43, GH51, CE15, CE16) (4), as well as pectinases (GH28, CE8) and also some CAZymes targeting fungal cell wall components (e.g. GH18 chitinases) (**Table 1**). Out of 23 identified AAs (18%), we found laccases (AA1) and class II peroxidases (AA2), LPMOs (AA9) and H_2_O_2_-generating enzymes (AA3, AA5). We estimated the abundance of the individual proteins using the Normalized Spectral Abundance Factor (NSAF) (**Dataset 1**). The mean %NSAF of all identified proteins was ∼ 0.3% and 52 proteins were above this value. The most abundant protein categories were CAZymes (52%), proteins of unknown function (35%) and proteases (8%). Among the most abundant CAZymes, we found the three cellobiohydrolases (GH7), three chitinases (GH18) and two laccases (AA1). Interestingly, three H_2_O_2_-generating enzymes (two AA3_2 and one AA5_1) were among these most abundant CAZymes (**Dataset 1**). Overall, the identity of the secreted proteins and their abundance were similar to those previously described for *P. cinnabarinus* or very closely-related species (26, 31–33). This validates our approach and indicates that the proteins preparation method and the oxidative treatment did not affect the retrieval of the secreted proteins, and that they were suitable for studying Met sensitivity to oxidation.

**Figure 1.**
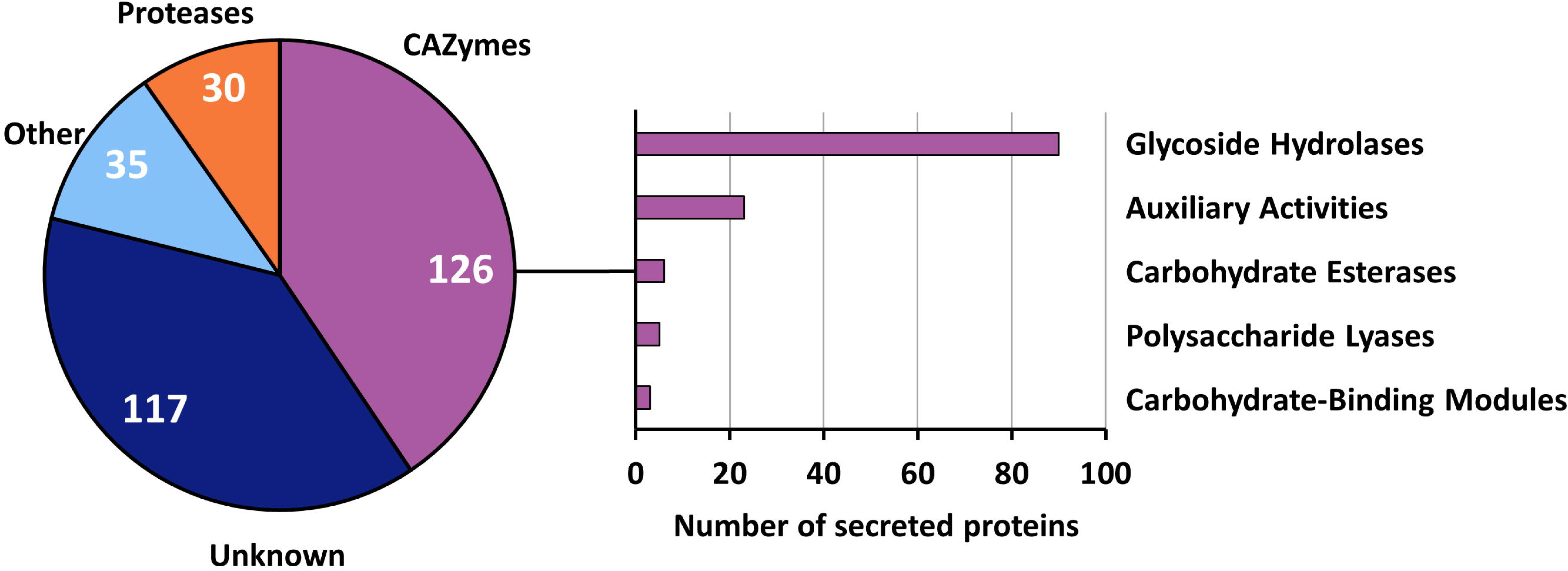
Distribution of the identified secreted proteins and CAZymes. Values shown are the aggregate of all replicates for each day (**Table 1; Dataset 1** (52)). Bar graphs represent the number of CAZymes per class.

**Table 1.**
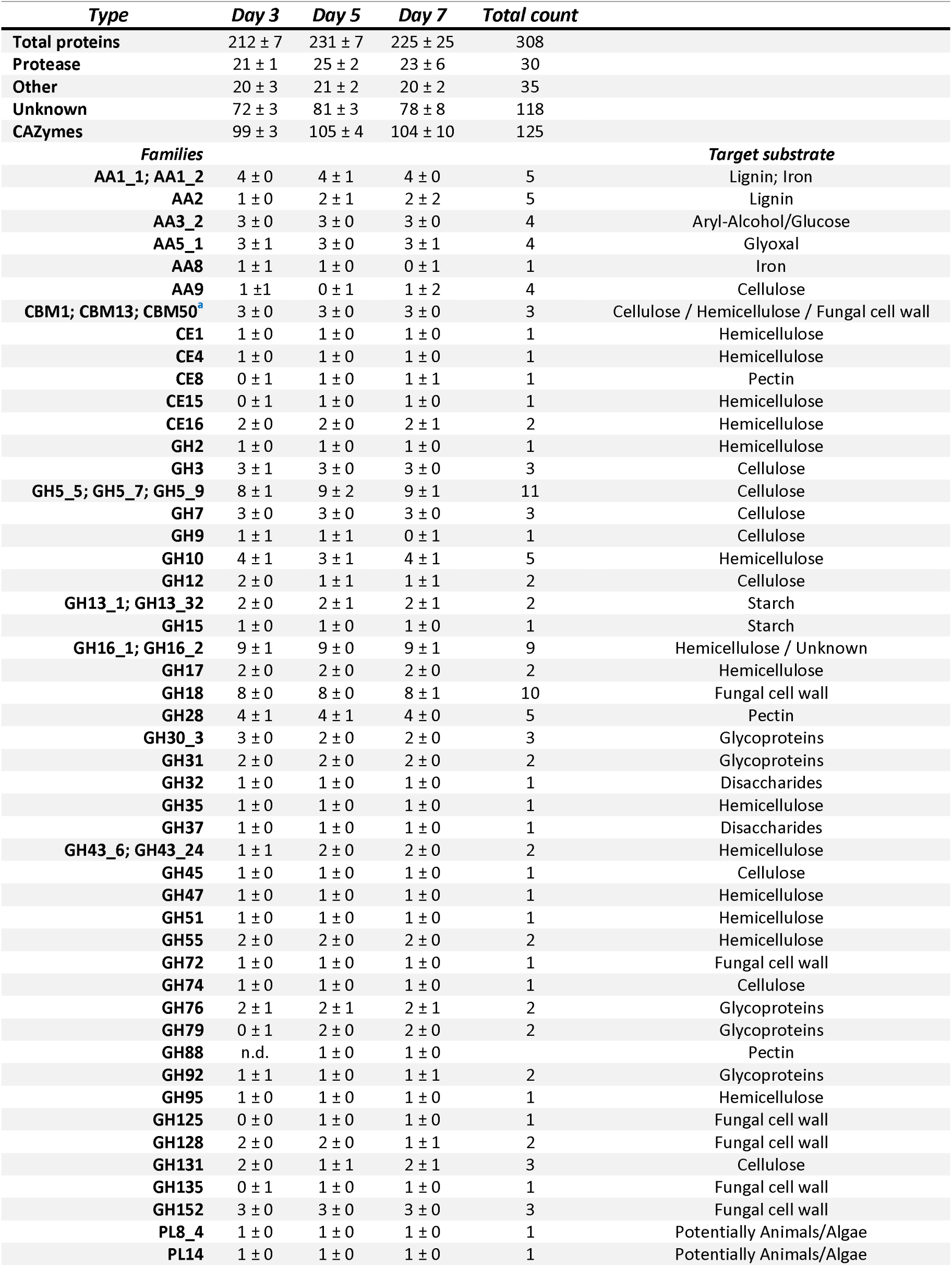
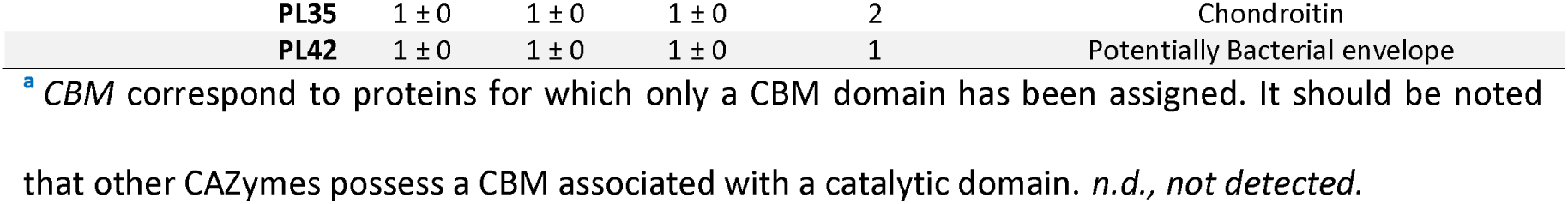
Number of identified secreted proteins. The list of individual proteins and their abundance is provided in the **Dataset 1** (52).

### Met content is reduced in fungal secreted proteins

First, we evaluated the Met content of the secreted proteins, which was ∼ 1.4% for each time-point (**Fig. 2A**). Then, to determine whether some abundant proteins with specific Met content could influence the overall content, we corrected the values by the proteins NSAF. The corrected Met contents were 1.21 ± 0.02, 1.33 ± 0.02, and 1.35 ± 0.02, for days 3, 5, and 7, respectively, without significant differences from the uncorrected values (**Fig. 2A**). These results indicate that the Met content in the proteins secreted by *P. cinnabarinus* is comprised between 1.2 to 1.4%. Strikingly, this value is twice lower than the general Met content of proteins, which is ∼ 2.4% in average for all protein sequences in the UniProt database (34). We investigated whether this characteristic may be applied to all proteins predicted to be secreted. *P. cinnabarinus* has 666 proteins predicted to be secreted with an average Met content of 1.55 ± 0.82%, while the non-secreted proteins have a higher mean of 2.07 ± 0.96% Met. (**Fig. 2B; Table S1; Dataset 2**). These results show that *P. cinnabarinus* secreted proteins contain significantly less Met than the non-secreted. Then, we compared the Met content of experimentally validated secreted proteins from fungi, metazoans, and plants with the Met content of intracellular proteins (**Fig. 2B; Table S1; Dataset 2**). The retrieved fungal secreted proteins were almost exclusively from Dikarya (Ascomycota and Basidiomycota) and have a mean Met content of 1.52 ± 0.74%, significantly lower than that of non-secreted proteins (2.17 ± 0.99%). This decrease in Met content for secreted proteins is conserved when Ascomycota and Basidiomycota are considered independently (**Fig. S2**). In contrast, the Met content of metazoans secreted proteins (2.68 ± 1.40%) is significantly higher than that of the non-secreted proteins (2.41 ± 1.07%). No statistical difference exists between the Met content in secreted (2.59 ± 1.55%) and non-secreted (2.55 ± 1.14%) proteins of plants (**Fig. 2B; Table S1; Dataset 2**). Altogether, these results show that the secreted proteins of fungi contain less Met than the intracellular proteins, and that this feature is not conserved in metazoans or plants.

**Figure 2.**
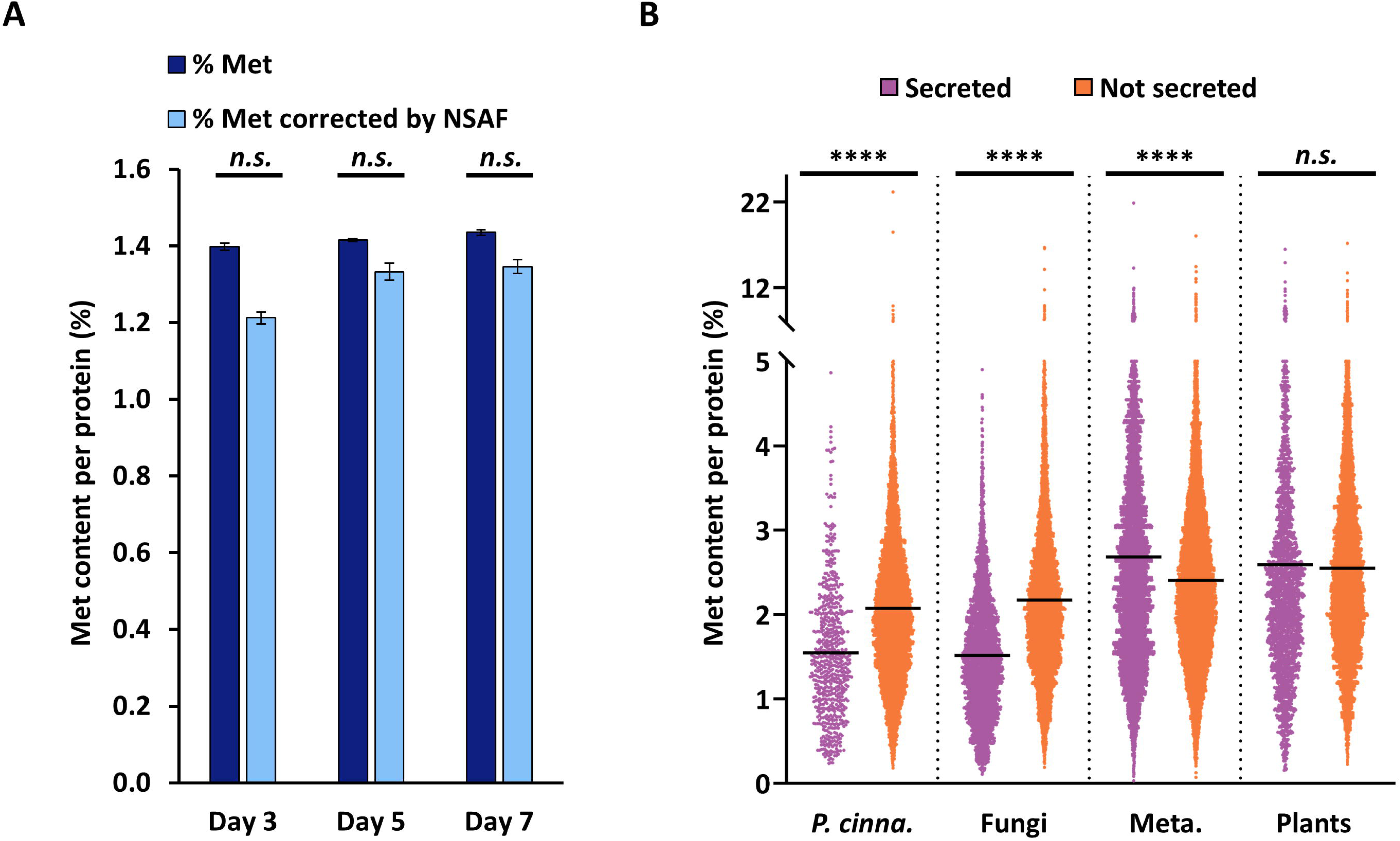
Met content of *P. cinnabarinus* secreted proteins and comparison with fungal, metazoan, and plant secreted and intracellular proteins. **A**) The Met percentage of identified *P. cinnabarinus* secreted proteins was calculated for each protein and averaged for the secretome of each day (*dark blue* bars). Alternatively, the Met percentage for each protein was weighted by the protein NASF and averaged (*light blue* bars). Values shown are the average of the 3 replicates ± standard deviation. **B**) Met percentages of all proteins predicted to be secreted or intracellular in the *P. cinnabarinus* were compared with the Met percentages of experimentally validated intracellular and secreted proteins from fungi, metazoans, and plants (**Table 1; Dataset 1** (52)). Mean percentage of Met are indicated with *black* lines. Statistical analysis was performed by unpaired t-test with Welch’s correction (\*\*\*\**P* ≤ 0.001; *n.s*., not significant).

### A monofunctional catalase protects secreted proteins from oxidation

The identification of oxidizable Met in *P. cinnabarinus* secreted proteins required accurate quantification of their oxidation levels. H_2_^18^O_2_ was used to prevent artifactual Met oxidation potentially occurring during sample preparation by blocking the Met reduced in the samples and converting them to Met sulfoxide or sulfone containing ^18^O atom (Met^18^O and Met^18^O_2_, respectively). The use of a concentrated H_2_^18^O_2_ solution (100 mM) should allow the oxidation of all the reduced Met and the reliable quantification of the initial levels of Met^16^O and Met^16^O_2_ (28, 29). For the day 3 secretome, the levels of Met carrying ^18^O or ^16^O, either as sulfoxide or sulfone, were ∼64% and ∼18%, respectively. Although ∼18% of Met in reduced form was still present, this indicated that most reduced Met in the culture conditions were blocked after the treatment with H_2_^18^O_2_. Of note, ^16^O or ^18^O sulfone levels reached ∼2% (**Fig. 3A; Table S2; Dataset 3**). On the contrary, in day 5 and day 7 secretomes, the levels of MetO and MetO carrying ^18^O were extremely low (∼1%), indicating that the blocking strategy failed for these samples (**Fig. 3A; Table S2; Dataset 3**). To evaluate whether this lack of oxidation by H_2_^18^O_2_ in secretomes of days 5 and 7 affected only Met residues, we reanalyzed our data by searching for two main forms of Pro carbonylation bearing ^18^O. For the day 3 secretome, we found slightly less than 4% of Pro oxidized with ^18^O. A similar content of ^16^O was found, indicating that the large majority of Pro were not oxidized (∼ 92%) (**Fig. 3B; Table S3; Dataset 4**). This low level of Pro oxidation was anticipated due to its low reactivity with H_2_^18^O_2_ (35). The percentage of ^18^O oxidation decreased to ∼0.5% in the day 5 and day 7 secretomes, while ^16^O oxidation levels remained comparable to those observed for the day 3 secretome. Thus, the trend for Pro is the same as for Met indicating that the lack of oxidation observed for Met was not specific to this residue. Proteins secreted during the days 5 and 7 of decay were not oxidized by exogenous H_2_^18^O_2_, unlike those secreted at day 3. These results prompted us to search for secreted enzymes that scavenge H_2_O_2_. A catalase stood out: almost absent at day 3 (0.02 ± 0.02 %NSAF), but more abundant at days 5 and 7, with %NSAF values of 0.33 ± 0.05 and 0.42 ± 0.01, respectively, above the mean of ∼ 0.3% found for all proteins (**Fig. 3C; Dataset 1**). When we measured the H_2_O_2_-consuming activity of the secretomes, we observed no activity at day 3, but strong activities at days 5 and 7 secretomes, in a perfect consistency with the catalase abundance (**Fig. 3C**). Although a few potential peroxidases and LPMOs were detected, their abundance was much lower than that of the catalase, and none followed a similar trend over the three days (**Table S4; Dataset 1**). These results indicated that this catalase was very likely responsible for most, if not all, of the H_2_O_2_-consuming activity. The sequence and modeled structure analyses revealed that this secreted catalase is a typical monofunctional catalase, similar to bacterial KatE enzymes or the peroxisomal enzyme found in animals and yeast cells (36) (**Fig. S3**). Altogether, these results show that, under our experimental conditions, *P. cinnabarinus* secreted a catalase at days 5 and 7, which, by consuming potential exogenous H_2_O_2_, protected the secreted proteins from oxidation, whereas at day 3, the secreted proteins are left without any protection against exogenous oxidant. Next, since Met oxidized with ^18^O were detected, we analyzed the nature of the oxidized proteins to identify Met residues with potential antioxidant or regulatory roles.

**Figure 3.**
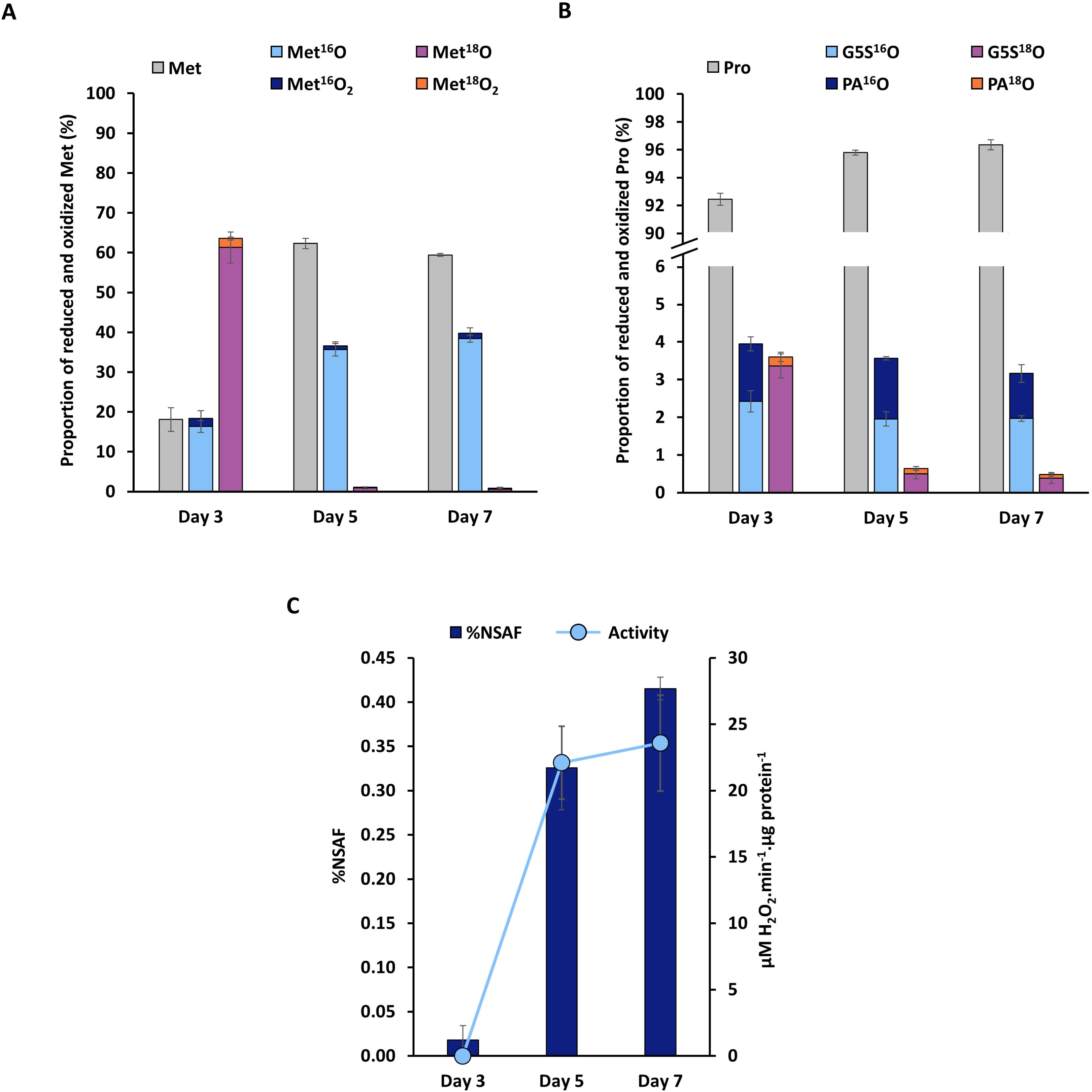
Levels of Met and Pro oxidation in the *P. cinnabarinus* secretomes and H_2_O_2_-consuming activity. **A**) Total levels of Met oxidation as sulfoxide and sulfone, either ^16^O or ^18^O calculated using the spectral counts of all Met-containing peptides and compared to the level of non-oxidized Met for each time point (**Table S2; Dataset 3** (52)). **B**) Total levels of Pro oxidation as glutamic 5-semialdehyde (G5S) and pyroglutamic acid (PA), either ^16^O or ^18^O calculated using the spectral counts of all Pro-containing peptides and compared to the level of non-oxidized Pro for each day (**Table S3; Dataset 4** (52)). **C**) %NSAF of the catalase A0A060STN2 (Dataset 1 (52)) and total H_2_O_2_-consuming activity at each time point.

### Most of the oxidations occur on a few secreted CAZymes

Among the 308 secreted proteins identified, we detected 265 Met belonging to 118 proteins (**Table S2; Dataset 3**). We selected only the Met-containing peptides present in all replicates for each secretome, with at least five spectral counts, in at least two out of the three replicates. At day 3, all selected Met were found to be oxidized with ^18^O, at least partially (**Fig. 4A; Dataset 3**). The total oxidation (^16^O and ^18^O) ranged from ∼ 20% to 100% for the 46 identified Met (**Fig. 4A**). Four Met, all from CAZymes, were fully oxidized and thus saturated with ^18^O: Met305 of a PL35, Met378 of a GH7, Met314 of a PL8, and Met239 of a CE16 (**Fig. 4A; Dataset 3**). Twenty-four proteins had oxidation levels ranging from ∼81% to 97% and 17 from 57% to ∼80%. Only the Met592 from an AA3_2 displayed a much lower value of total oxidation (∼20%) (**Fig. 4A; Dataset 3**). The ^16^O-oxidation levels were confidently evaluated for the four ^18^O-saturated Met and should correspond to the levels “naturally” found during the wood decay. However, for all other Met with an oxidation value less than 100%, the saturation with ^18^O was incomplete and some artifactual oxidation may have occurred during the sample preparation, and thus the level of ^16^O might be overestimated. For these proteins, we assumed that the percentage of ^16^O found during wood decay was necessarily equal to or lower than the measured one. The percentage of oxidation as ^16^O ranged from 0% to 47% with a mean of 18% (**Fig. 4A; Dataset 3**). This value is consistent with the mean percentage of Met^16^O oxidation calculated considering all Met containing-peptides found (**see Fig. 3A**) and indicated that most of the oxidation was concentrated on these few secreted proteins. Concerning day 5 and day 7 secretomes, the very low levels of ^18^O oxidation precluded the use of the blocking strategy. However, to find proteins with Met^18^O may allow to consider H_2_^18^O_2_ as a probe for highly reactive residues, because of the low H_2_^18^O_2_ concentration they were exposed to, due to the catalase activity. We found eight and nine Met bearing ^18^O in the proteins secreted at day 5 and day 7, respectively, and five were common to both samples (**Fig. 4B; Dataset 3**). The percentages of ^18^O ranged from 2% to ∼14%, and from ∼1% to ∼23%, for the secreted proteins of day 5 and day 7, respectively. Among the 11 oxidized Met, only three were uniquely found at day 5 or day 7: Met533 of a GH95 at day 5, and Met284 and Met160 belonging to a PL35 and a GH3, respectively, at day 7 (**Fig. 4B; Dataset 3**). This indicates that most of the proteins found oxidized with ^18^O at day 5 or 7 were also oxidized at day 3, highlighting their high reactivity to H_2_O_2_. We found a total number of 49 oxidizable Met belonging to 28 proteins, representing ∼24% and ∼19% of all the identified Met and proteins from the secretome, respectively (**Fig. 4C; Dataset 3**). Interestingly, the 18 identified CAZymes represented most of them (64%), while proteases and proteins of unknown function both represented 18% with five identified proteins. The most represented CAZymes were GHs with 10 identified proteins. These results indicates that the proportion of oxidized CAZymes was enriched compared to the different types of proteins present in the secretomes (**compare Fig. 4C with Fig. 1**).

**Figure 4.**
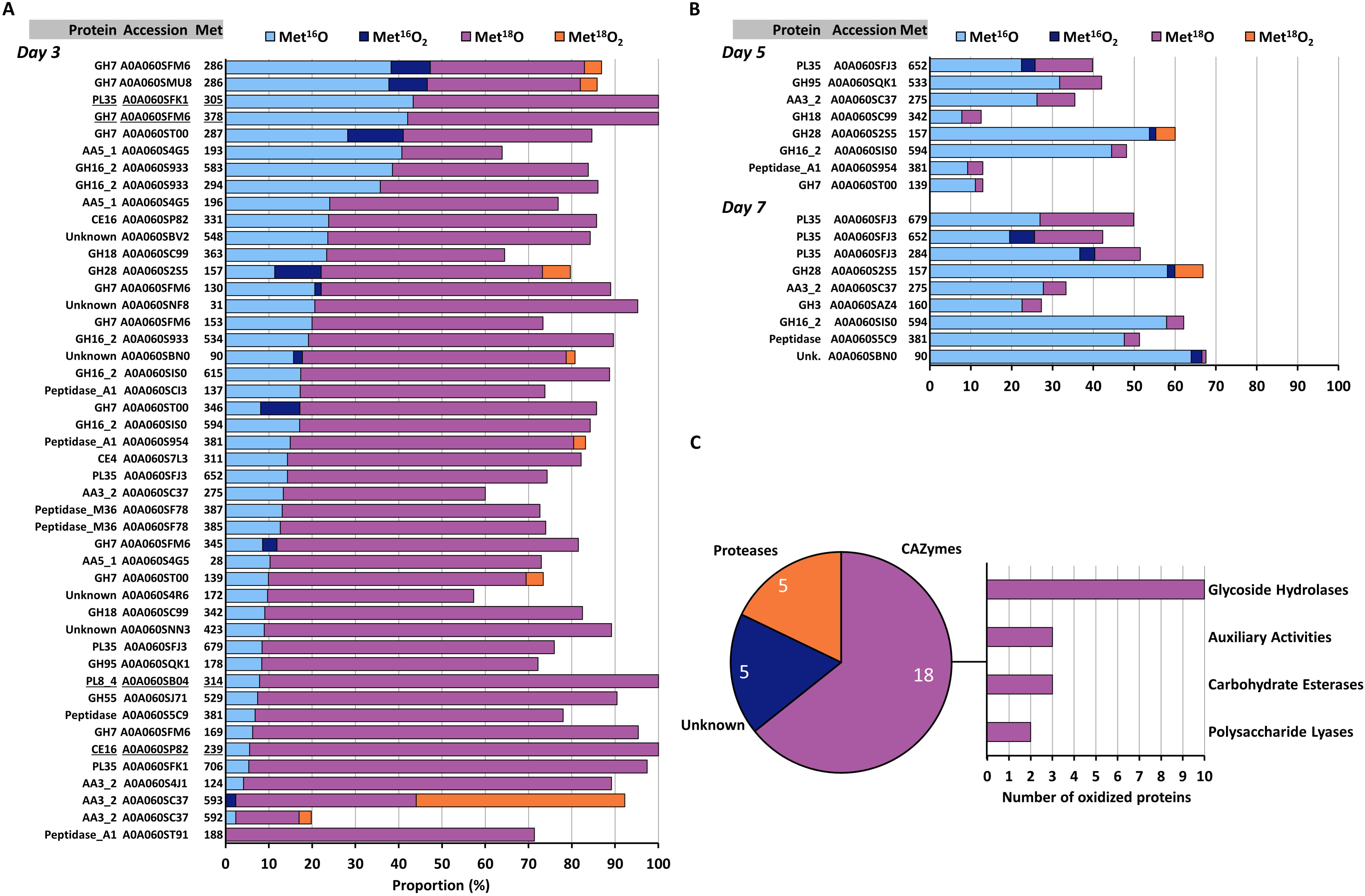
Identity of oxidized Met and their levels of oxidation in *P. cinnabarinus* secretomes. **A**) Proportion of the different forms of oxidation for the Met identified in the secreted proteins at day 3. Proteins are ranked based on the percentage of ^16^O (MetO and MetO). The four Met fully oxidized are underlined. Values shown are the mean of oxidation without standard deviation, omitted for clarity. All values are in **Dataset 3** (52). **B**) Proportion of the different forms of oxidation for the Met identified in the secreted proteins at day 5 and day 7. Proteins are ranked based on the percentage of ^18^O (MetO and MetO). All values are in **Dataset 3** (52). **C**) Distribution of the identified secreted proteins and CAZymes with oxidized Met. Values shown are the aggregate of all replicates for each time point (**Dataset 3** (52)). Bar graphs represent the number of CAZymes per class.

### Oxidation occurs in homologous proteins, at multiple sites and in hotspots

Interestingly, among the five most oxidized, three Met belong to the three GH7 (A0A060SFM6, A0A060SMU8, A0A060ST00, named A, B, and C, respectively), at position 286 of the GH7 A and B, and at position 287 of the GH7 C (**Fig. 4A**). A sequence alignment showed that these positions are conserved between these GH7 homologs (**Fig. 5A**). These Met were oxidized with similar levels of Met^16^O (between 28% and 38%) and Met^16^O_2_ (between 9% and 13%). We also detected seven other oxidized Met on these GH7s, four were uniquely on the GH7 A, one on the GH7 B, and two were in a position conserved in GH7 A and B. Since we had assessed the overall level of Pro oxidation, we quantified the percentage of oxidation of individual Pro, considering only those labeled with ^18^O (**Dataset 4**). GH7 A had four oxidized Pro, two of which were in positions conserved into the GH7 B (**Fig. 5A**). Interestingly, we found that oxidation of Met and/or Pro occurred on homologs of GH16_2, PL35, proteases and GH152 (**Fig. 4; Fig. S4**). This represented 11 proteins among the 33 proteins identified having oxidized residues (28 with Met with or without Pro and 5 with Pro only) (**Fig. 4; Fig. S4; Datasets 3; 4**). Moreover, 19 out of the 33 identified proteins were oxidized on multiple residues, from two to 10. A GH152 was oxidized on 8 Pro, while the GH7 A was the protein with the highest number of oxidation sites (**10), including six Met and four Pro (Fig. 4; Fig. 5; Fig. S4; Datasets 3; 4**). Finally, several oxidation sites were located close to each other in the protein sequences, forming “hotspots” like the Pro343 and Met345, and three other Pro of the GH7 A (**Fig. 5A**). In total, we counted 24 occurrences of sites containing two oxidized residues in the protein sequences (**Fig. 5B**). Analysis of a tridimensional model of GH7 A revealed that oxidized residues distant on the sequence may be close on the structure, as the Met130 and Met160, the Met153 in the vicinity of three Pro, and the Met286 clustered with the Pro343 and Met345 (**Fig. 5C**). Altogether, these results indicate that most of the oxidation affects a relatively small set of CAZymes that can be oxidized at multiple positions, with some hotspots. Moreover, the fact that oxidized residues were found at similar positions within the homologs of *P. cinnabarinus* secreted proteins suggests that these residues have been conserved to fulfill antioxidant or regulatory roles.

**Figure 5.**
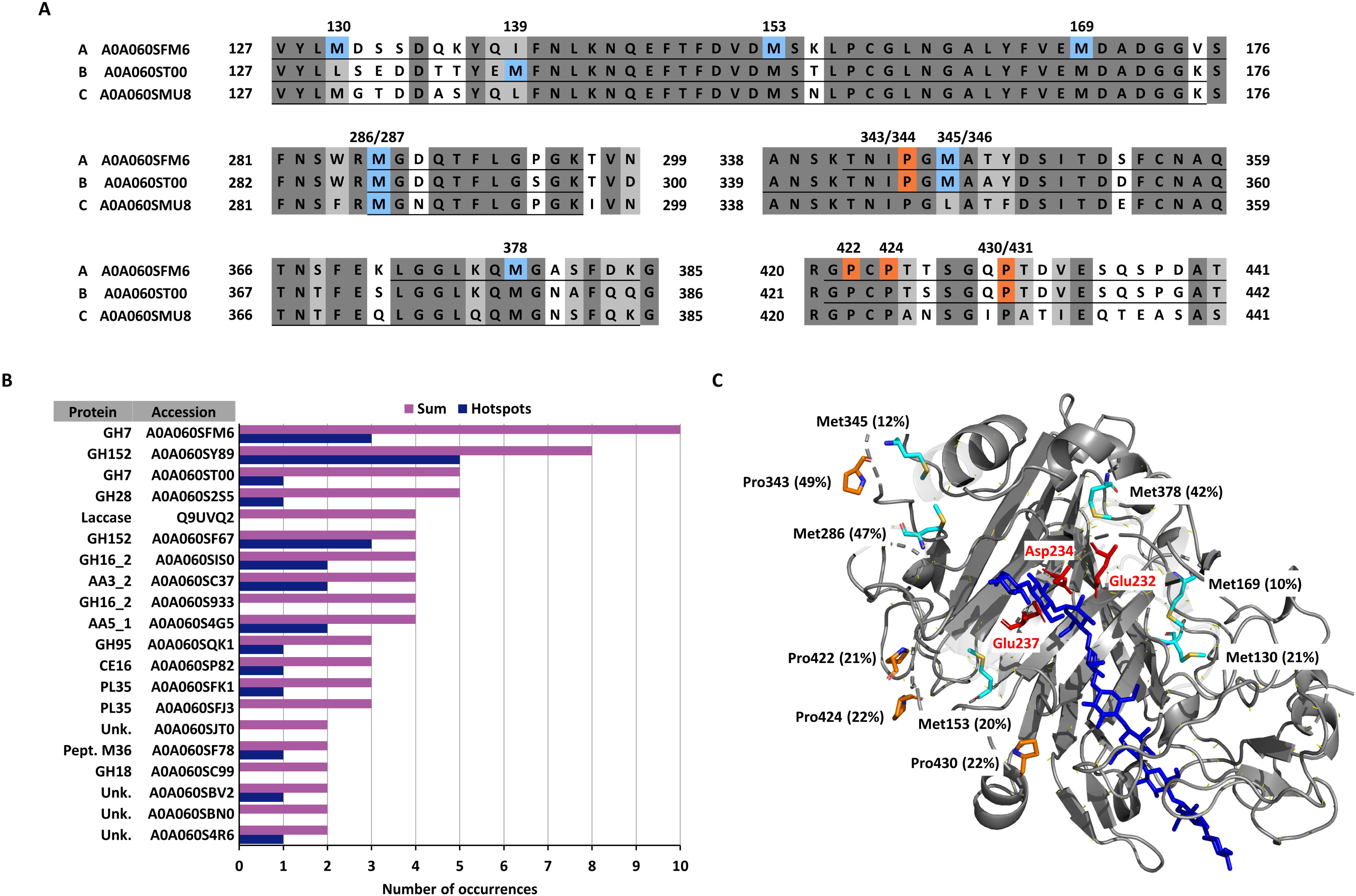
Partial sequence alignment of GH7, numbers of oxidation sites and hotspots in identified secreted proteins and tridimensional model of GH7. **A**) Oxidized Met and Pro residues are on *blue* and *orange* background, respectively. *Dark* and *light grey* backgrounds represent strictly conserved and similar amino acids, respectively. Position and percentage of oxidation of the residues are indicated above the sequence. For Pro, the percentage presented correspond to the sum of all oxidized forms, either ^16^O or ^18^O at day 3, excepted the one indicated with ‘D5’, corresponding to Pro oxidation at day 5. Underlined sequences represent peptides detected by LC-MS/MS. Accession numbers are on the left of the sequences. **B**) Values correspond to the total number of oxidizable Met and Pro sites found per protein. Hotspots correspond to the number of times that two oxidizable sites were located within a window of ten amino acids on the protein sequence. All data are presented in **Datasets 3 and 4** (52). **C**) 3D model of the GH7 A0A060SFM6 downloaded from Alphafold database (77). The position of the cellonoanose substrate (in *blue*) was modelized using the experimentally determined structure of *Trichoderma reesei* Cel7A (PDB #4C4C). Met and Pro found oxidized are represented as *cyan* and *orange* sticks, respectively. Catalytic residues are represented as *red* sticks. Percentage of oxidation at day 3 are in brackets. Some parts of the protein were rendered transparent for clarity. Structure manipulation and image generation were done using PyMol 2.3.2 (https://pymol.org/2/).

### Conservation study suggests a list of residues with antioxidant or regulatory roles

To determine whether the 89 oxidized residues found in the 33 *P. cinnabarinus* proteins were conserved in fungi, we searched for fungal homologs of the proteins and quantified the conservation of the oxidized residues. Our hypothesis was that oxidizable residues playing antioxidant or regulatory roles are more likely to be conserved among distantly related organisms, as shown for Met of actin or of the ffh/SRP proteins (37–39). To avoid overrepresentation of sequences coming from closely related organisms, the search was made in 120 representative fungal species, covering the nine fungal phyla (**Dataset 5**). Overall, the conservation of Met or Pro ranged from 1% to 100%, with a mean of 49%. Considering only residues with conservation values above 75%, and found in more than 80 sequences, we obtained a list of nine Met and nine Pro at conserved positions in 14 secreted proteins (**Table 2**). The GH7 A has four conserved Met, among which Met 169 conserved in 99% of the homologs, and a conserved Pro. Other Met are conserved in the GH7 B and C, and in two GH16_2, one GH18, one AA3_2 and a peptidase M36, present in all fungi (**Table 2; Dataset 5**). Regarding Pro, three are conserved in all homologs, at positions 614, 143 and 441 of a GH16_2, a GH28 and a laccase, respectively. Conserved Pro were also found in a AA3_2, an unknown protein and a GH152 (**Table 2; Dataset 5**). These results allowed identification of proteins with oxidizable Met and Pro residues strongly conserved across the fungal phyla.

**Table 2.**
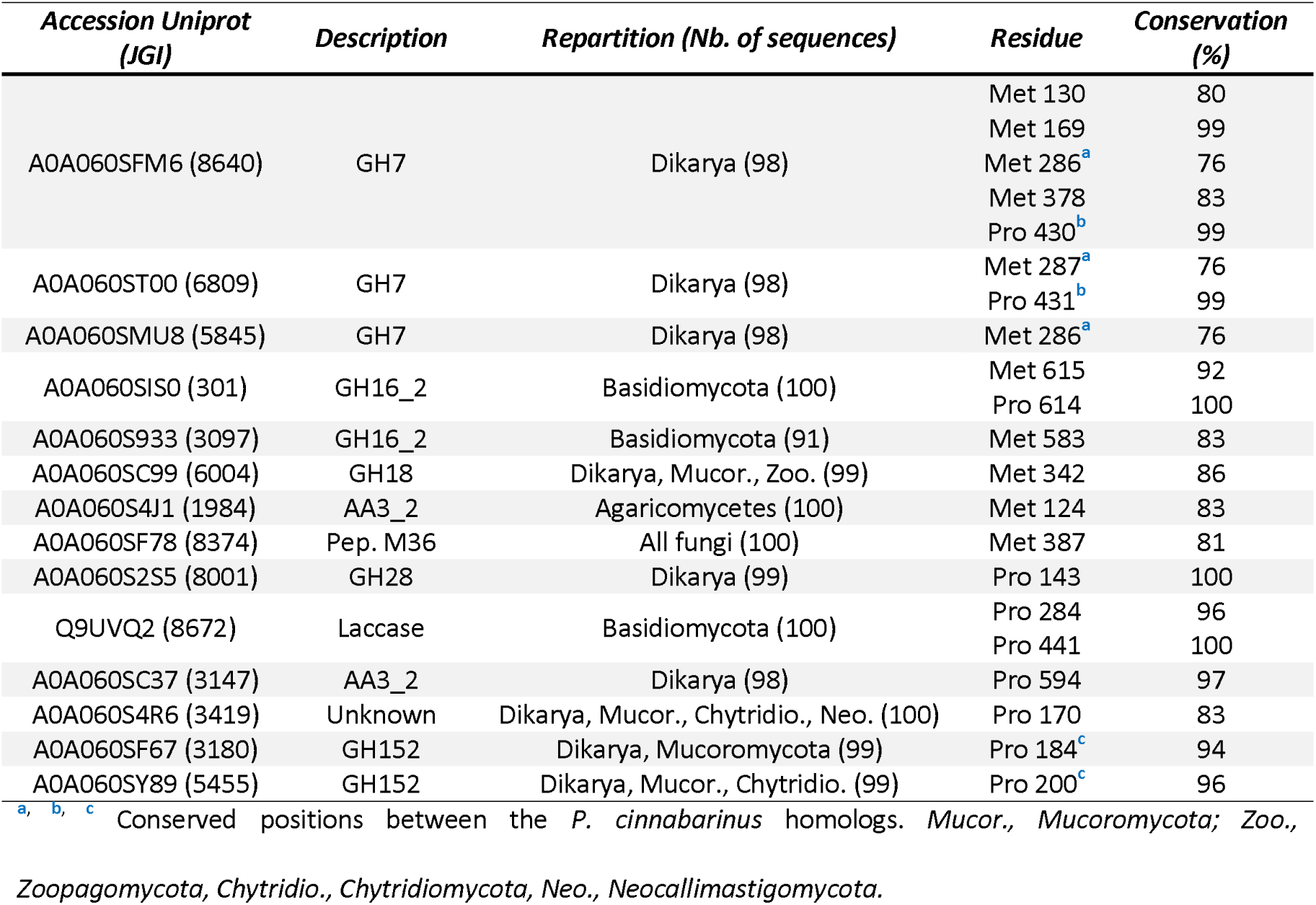
*P. cinnabarinus* secreted proteins with oxidized residues conserved across fungal proteins. Only proteins for which the number of identified homolog sequences identified was higher than 80 were considered. The complete list of identified secreted proteins is presented in **Dataset 5** (52).

## Discussion

Taking a white-rot wood decayer as model, we showed that fungal secreted proteins have a strongly reduced Met content (∼1.4%) compared to the non-secreted (∼2.1%) and that this feature is not conserved in metazoans and plants. This reduced Met content could contribute to the tolerance of the secreted enzymes to oxidation by reducing the number of sensitive residues. This could explain, at least in part, the strong resistance of some CAZymes activities exposed to high concentrations of Fenton-generated ROS observed in the brown-rot *Postia placenta* (16, 40). Interestingly, these data contrast with those obtained from animal studies which showed that proteins encoded by the mitochondrial genome may contain up to 6-10% Met. This high content could act as an endogenous antioxidant with the exposed Met scavenging ROS which are then removed by methionine sulfoxide reductases in a cycle of oxidation and reduction (41). A similar mechanism was proposed in bacteria (42, 43). The absence of external system of MetO reduction precludes the use of such mechanism with fungal secreted proteins (24). The protection of the secreted proteins from oxidation is most likely also due to the activity of the secreted catalase we identified (**Fig. 3; Table S4**), consistent with previous suggestions showing that the addition of catalase to CAZymes cocktails reduces inactivation over time (44, 45). The secretion of a catalase by *P. cinnabarinus* was unexpected, as it has not been previously detected in secretome analysis (26, 31). Moreover, catalase extracellular activity was also not measured for the white-rot *Phanerochaete chrysosporium* (46). However, searching for secreted catalase homologs, we found that it is widespread in Basidiomycota, and particularly in white-rot fungi (**Table S5**). This conclusion contrasts with a previous phylogenetic analysis, which argued that Basidiomycota possess only monofunctional catalases not predicted to be secreted because of the lack of a signal peptide (47). Lack of detection of the protein or its activity may be due to specific expression conditions. In white-rot, it is surprising that a catalase was expressed along with H_2_O_2_ producing and consuming enzymes. However, catalases have Km values in the mM range (44), and their action could result in lowering the H_2_O_2_ concentration below a level that is damaging to proteins, but allows the activity of H_2_O_2_-driven oxidoreductases.

Using H_2_^18^O_2_-based proteomics, we identified oxidizable Met and highlighted that CAZymes were the most oxidized secreted proteins during wood-decay, and that Met oxidation could occur on several residues, sometimes clustered in hotspots. Clustering was also observed in bacterial proteins (48), suggesting a universal phenomenon. The blocking strategy we used is based on a saturation of reduced Met with ^18^O. However, saturation was difficult to achieve, with only 4 Met saturated with ^18^O in the day 3 secretome (**Fig. 4**). An initial denaturation step would have probably allowed H_2_^18^O_2_ to access the Met buried in the protein core (28). However, as our goal was to identify secreted proteins with potential oxidizable Met we chose not to denature them. Our idea was that an oxidizable Met should be accessible to an oxidant, either natural or H_2_^18^O_2_. Indeed, the results at day 3 argue for the accessibility of Met since all identified Met but one carried both ^16^O– and ^18^O-oxidation, with 43 Met out of 46 for which the percentage of ^18^O was higher than that of ^16^O. If a Met would not have been accessible to an oxidant in the folded protein, the H_2_^18^O_2_ would not have oxidized it, and we could have observed ^16^O, very likely coming from artifactual oxidation, but no ^18^O. However, as artefactual ^16^O-oxidation cannot be totally ruled out, and potentially happened on the remaining non-oxidized fraction of a considered Met during the proteomics procedure, we considered that the “natural” percentage of ^16^O was necessarily lower or equal to those measured. In our study, the mean percentage of ^16^O was 18%, while in studies on animals using similar approaches, it was ∼4% (27, 28), arguing for potential artifactual oxidation. However, while intracellular proteins are protected by methionine sulfoxide reductases and ROS scavengers, the MetO in secreted proteins cannot be reduced, and no H_2_O_2_-consuming was detected in the secretome at day 3. Thus, to find high levels of MetO in fungal secreted proteins is expected. Interestingly, nine Met were oxidized as ^16^O sulfone among which five were also oxidized as Met^18^O (**Fig. 4**), confirming their propensity to form sulfone during wood decomposition.

Some of the Met we found oxidized in P. cinnabarinus are conserved throughout the fungal kingdom, suggesting potentially important roles (**Table 5**). They could act as antioxidants, as for the GH7 whose Met and Pro are located at the surface of the proteins and in the substrate cleft (**Fig. 5C**), potentially preventing critical residues from being oxidized, as proposed for a glutamine synthetase (23). Met sensitive to oxidation could also have a functional role altering enzyme activity like for a GH18 in which a conserved oxidized Met is located near the catalytic residue (**Fig. S3**). This type of regulation was shown for a GH27 galactosidase of *T. reesei*, for which Met oxidation increased activity (25). In conclusion, our findings highlight a common mechanism in fungi preventing oxidation of secreted proteins and, pinpoint redox-regulated CAZymes to be further characterized.

## Material and methods

### *Culture of* P. cinnabarinus *on aspen wood*

The dikaryotic strain *Pycnoporus cinnabarinus* CIRM-BRFM 145 was grown for 3, 5 or 7 days at as described (26). The media contained maltose (2.5 g.l^−1^) and *Populus tremuloides* aspen sawdust particles of size < 21⍰mm (15 g. l^−1^). Six replicates were mixed in pairs to obtain three replicate samples per time-point.

### *Preparation of* P. cinnabarinus secretomes and protein oxidation with H ^18^O

The culture supernatants containing the secreted proteins were centrifugated (1 h, 30,000xg, 10 °C) and sterilized by filtration (0.22 µm; Express Plus; Merck Millipore). The supernatants were equilibrated to pH 7 with 0.5 mM NaOH and the proteins were purified using two ion exchange chromatographies (**Fig. S1**): first, an anion exchange on DEAE Sephadex A-25 (Cytiva) and a cation exchange on CM Sephadex A-25 (Cytiva) for proteins not retained on the anion exchanger. Both resins were initially equilibrated with 20 mM sodium phosphate buffer (pH 7). Elution of the secreted proteins was performed with 1 M NaCl. Fractions from both anion and cation exchangers were pooled, desalted on Sephadex G-25 (PD-10 Desalting columns, Cytiva) in 50 mM sodium acetate (pH 5.2) and concentrated using Vivaspin polyethersulfone membrane (3 kDa cut-off, Sartorius). Protein concentrations were determined using the Bradford method (Bio-Rad) and the secretomes were used immediately. Secreted proteins (50 µg) were incubated 30 min with 1 mM hydroxylamine, desalted on Sephadex G-25 (PD SpinTrap, Cytiva), then incubated with 100 mM of H_2_^18^O_2_ (Sigma-Aldrich) for 1 h in the dark, then desalted. Initial concentration of commercial H_2_^18^O_2_ was estimated to be ∼ 450 mM using FOX method (49). Protein concentrations were determined, and the samples were stored at –20°C.

### Proteomics analysis

Proteins (15 µg) were precipitated by trichloroacetic acid and processed for trypsin proteolysis as described (50). Peptides (5 µl) were injected into a nanoscale Easy-Spray™ PepMap™ Neo 2 µm C18 (75 µm x 750 mm) capillary column (Thermo Scientific) operated with a Vanquish Neo UHPLC system (Thermo Scientific) and resolved with a 65-min gradient of CH_3_CN (5-40%), 0.1% formic acid, at a flow rate of 0.25 µL per min. Data-dependent acquisition analysis of the peptides was performed with an Orbitrap Exploris 480 tandem mass spectrometer (Thermo) as described (51). Each full scan of peptide ions in the high-field Orbitrap analyzer was acquired from m/z 350 to 1800 at a resolution of 120,000, selecting only 2+ and 3+ charged precursors for high-energy collisional dissociation and with a dynamic exclusion of 10 seconds. MS/MS spectra of fragmented precursors were acquired at a resolution of 15,000 and assigned to peptide sequences by the MASCOT Server 2.5.1 search engine (Matrix Science) using the *P. cinnabarinus* annotated proteome, with fixed and variable modifications. Data are available in **Datasets 3; 4** of associated preprint (52). Proteins were validated when at least two different peptides (p-value <0.01) were detected, resulting in a protein identification false discovery rate below 1%. Protein abundance was estimated using the Normalized Spectral Abundance Factor (NSAF) (53). The Met content of the proteins was calculated using the entire sequence. The Met content corrected by the NSAF was calculated using the following formula:

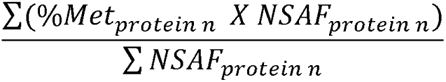

With %*Met_protein n_* and *NSAF_protein n_* corresponding to the percentage of Met and the NSAF value of the individual proteins identified, respectively. The percentage of modification of Met or Pro residues was calculated by dividing the number of spectral counts (SC) of the residue with the modification by the total number of SC for the residue and then multiplying the result by 100. The mass spectrometry proteomics data have been deposited to the ProteomeXchange Consortium via the PRIDE (54) partner repository with the dataset identifier PXD046271 and 10.6019/PXD046271.

### H_2_O_2_-consuming activity

Secreted proteins (20 µg) purified from day 3, 5 or 7 cultures as described before were incubated with H_2_O_2_ (30 mM) for 1 h. Then, the proteins were removed from the samples by ultrafiltration using Vivaspin 3 kDa cut-off and the filtrate was transferred into 1 ml quartz cuvette. The residual H_2_O_2_ concentration in the filtrate was titrated by measuring the absorbance at 240 nm using a molar extinction coefficient value of 43.6 M^-1^.cm^-1^ for H_2_O_2_. For each a day a control was made incubating the H_2_O_2_ in the protein purification buffer without proteins. The experiment was performed with a biological triplicate, and a technical duplicate. The activity of consumption was calculated as follows:

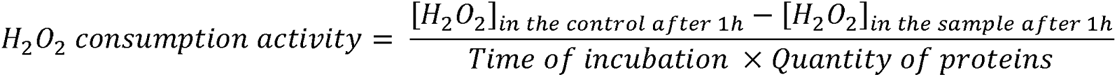

### *Analysis of Met content in* P. cinnabarinus, *fungi, metazoans, and plants proteins*

*P. cinnabarinus* proteins were predicted secreted if they fulfilled three conditions: (i) presence of a secretion signal peptide, (ii) absence of endoplasmic reticulum retention motif and (iii) absence of transmembrane helix outside the signal peptide, as described (26). Experimentally validated fungal secreted proteins were retrieved from proteomics analyses (7, 26, 31–33, 55–65), from Uniprot (www.uniprot.org) and FunSecKB2 (66). Metazoans experimentally validated secreted were retrieved from Uniprot, MetazSecKB (67) and the Human Protein Atlas (68, 69). Experimentally validated plant secreted proteins were retrieved from Uniprot, PlantSecKB (70) and the SUBA database (71). The datasets of fungal, metazoans and plants non-secreted proteins were made using proteins retrieved from Uniprot, the Human Protein Atlas and the SUBA database. For all datasets, only manually curated proteins, and those found with least 5 SC in proteomics, were conserved. Membrane proteins, proteins shorter than 50 amino acids, or not starting with Met were discarded. Statistical analyses were unpaired t-test with Welch’s correction. The numbers of proteins for each category are described in **Table S1** and the protein sequences are available in **Datasets 2** (52).

### Conservation of proteins and oxidized residues

A hundred and twenty fungal genomes were selected based on their public availability in the MycoCosm database (72) and their representativity in the 9 fungal phyla (73, 74). Targets of the oxidized proteins were searched using BLASTP (v 2.6.0+) (75). The 100 sequences with the highest e-values were aligned using MAFFT tool (version 6.864b) (76). The conservation of individual residues was made by counting its occurrence in all the sequences. Data are available in **Datasets 5** (52).

## Acknowledgments

The authors kindly thank the CIRM-CF for providing the *P cinnabarinus* CIRM-BRFM145 strain, Dan Cullen (Forest Product Laboratory, USDA, Madison, WI, USA) for providing *Populus tremuloides* wood, and Marie-Noëlle Rosso (Aix Marseille Univ., INRAE, Biodiversité et Biotechnologies Fongiques) for critical reading of the manuscript. This work received support from the French government under the France 2030 investment plan, as part of the Initiative d’Excellence d’Aix-Marseille Université —IM2B (AMX-19-IET-006) for L.M. and L.T. – A*MIDEX (AMX-21-PEP-028) for L.T. The authors would like to thank the NovoNordisk foundation (OxyMiST project, grant number NNF20OC0059697) and the Commission Nationale des Outils Collectifs (CNOC) of INRAE for funding.

## CRediT author statement

**Lise Molinelli**: Conceptualization, Methodology, Validation, Formal analysis, Investigation, Data Curation, Writing – Original Draft, Writing – Review & Editing, Visualization, **Elodie Drula**: Software, Methodology, Formal analysis, Investigation, Data Curation, **Jean-Charles Gaillard**: Formal analysis, Investigation, **David Navarro**: Investigation, Resources, **Jean Armengaud**: Formal analysis, Resources, Data Curation, Writing – Review & Editing, **Jean-Guy Berrin**: Resources, Writing – Review & Editing, Funding acquisition, **Thierry Tron**: Conceptualization, Resources, Writing – Review & Editing, Supervision, Funding acquisition, **Lionel Tarrago**: Conceptualization, Methodology, Software, Validation, Formal analysis, Investigation, Resources, Data Curation, Writing – Original Draft, Writing – Review & Editing, Visualization, Supervision, Funding acquisition.

## Notes

### Competing Interest Statement

The authors have declared no competing interest.

### Summary of Updates

– Figure 2 revised – Methods updated – Results description improved – Discussion improved – Dataset 2 modified – Supplementary data modified

