## Supplementary data for "Methionine oxidation of Carbohydrate-Active enZymes during white-rot wood decay"

**SUPPLEMENTARY INFORMATION**

**List of supplementary data**

| Figure S1. Secretome preparation and protein oxidation. | Page 2 |
| --- | --- |
| Figure S2. Met content of Ascomycota and Basidiomycota secreted proteins and comparison with intracellular proteins.  *Related to Figure 2* | Page 3 |
| Figure S3. Sequence and structure alignments of the *P. cinnabarinus* secreted catalase.  *Related to Figure 3* | Page 4 |
| Figure S4. Partial sequence alignment of GH16_2 (A), PL35 (B), protease (C) and GH152 (D) with identified oxidized residues.  *Related to Figure 5* | Page 5 |
| Figure S5. 3D model of GH18 (B) highlighting oxidized and conserved Met. | Page 6 |
| Table S1. Met content in secreted and non-secreted proteins of *P. cinnabarinus*, fungi, metazoans, and plants. | Page 7 |
| *Related to Figure 2* |  |
| Table S2. Levels of Met oxidation in secretomes obtained at day 3, 5 and 7. | Page 8 |
| *Related to Figure 3* |  |
| Table S3. Levels of Pro oxidation in secretomes obtained at day 3, 5 and 7. | Page 8 |
| *Related to Figure 3* |  |
| Table S4. Abundance of detected H_2_O_2_-consuming enzymes in the secretomes obtained at day 3, 5 and 7.  *Related to Figure 3* | Page 9 |
| Table S5. Number of predicted secreted catalases in fungi. | Page 9 |

**List of additional datasets**

| Dataset 1. List of identified secreted proteins and their spectral counts. |
| --- |
| Dataset 2. List of predicted secreted and non-secreted proteins of *P. cinnabarinus* and experimentally validated secreted and non-secreted proteins from fungi, metazoans and plants. |
| Dataset 3. Percentage of the different forms of Met for the identified proteins. |
| Dataset 4. Percentage of the different forms of Pro for the identified proteins. |
| Dataset 5. Conservation of secreted proteins and their oxidized residues in a representative set of fungal species. |

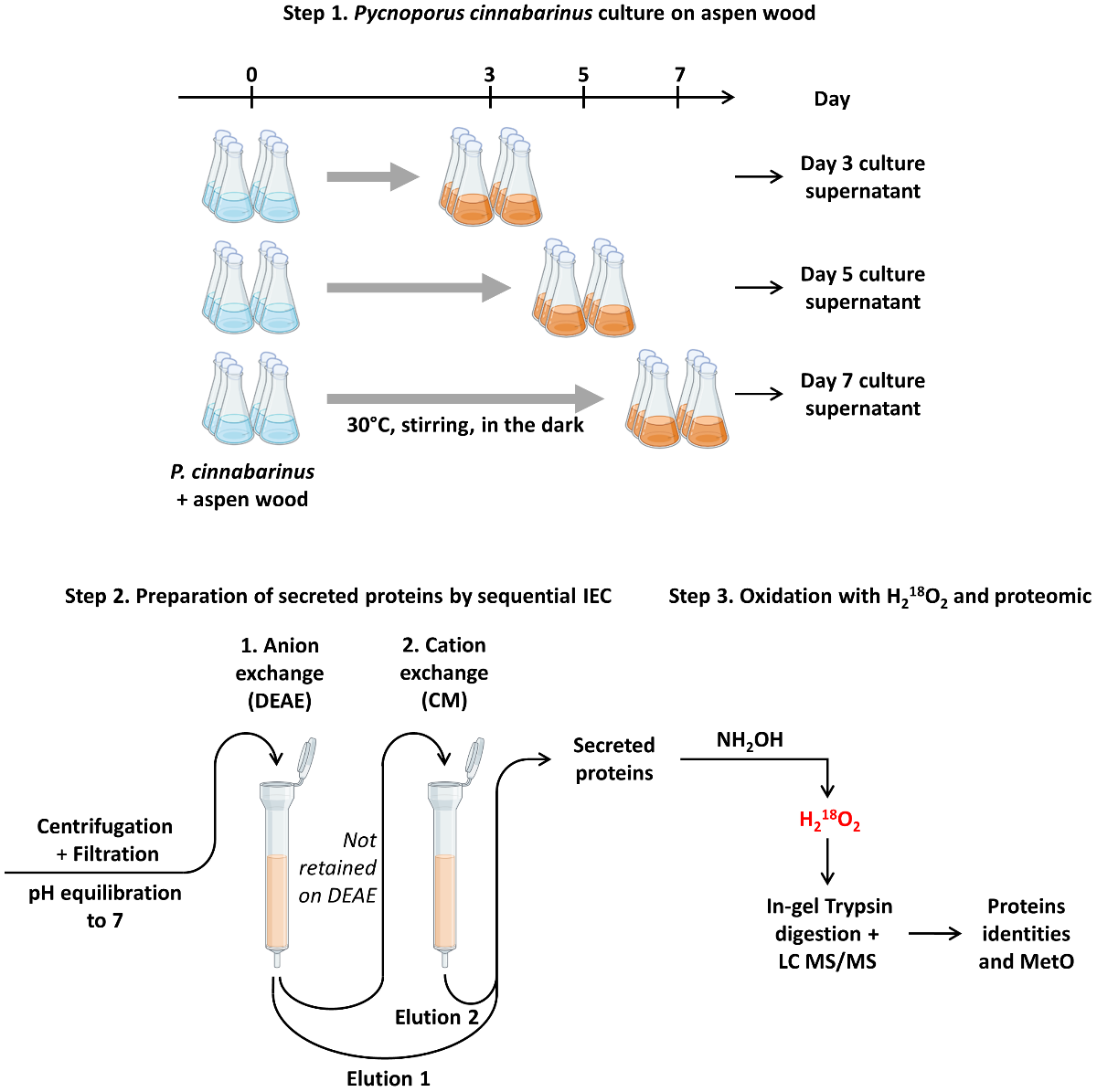

**Figure S1. Secretome preparation and protein oxidation.** The preparation of *P. cinnabarinus* secretomes and the oxidation of the secreted proteins were performed in 3 steps: *Step 1.* *P. cinnabarinus* was cultivated in the presence of *Populus tremuloides* sawdust (particle size < 2 mm) (15 g.L^-1^). Cultures were initiated in 18 flasks and 6 were used to prepare the secretomes of time point (day 3, 5 or 7). For each time point, the supernatants of 2 flasks were pooled to obtain one of the 3 replicates. *Step 2.* The supernatants were centrifuged, filtered and the pH adjusted to 7 with NaOH. The supernatant was then loaded onto an anion exchange column (DEAE). The solution containing the proteins not bound to the column during loading nor washing was loaded onto a cation exchange column (CM). The proteins retained on each column were eluted with NaCl (1M), pooled, desalted in 50 mM sodium acetate (pH 5.2) and concentrated by ultrafiltration (3 kDa cut-off). *Step 3*. The proteins (70 µg) were incubated with 1 mM hydroxylamine (NH_2_OH) to inhibit H_2_O_2_-consumming activity, desalted, and incubated with of H_2_^18^O_2_ (100 mM, 1 h). After desalting, 10 µg were used for in-gel trypsin digestion and peptide identification by LC-MS/MS.

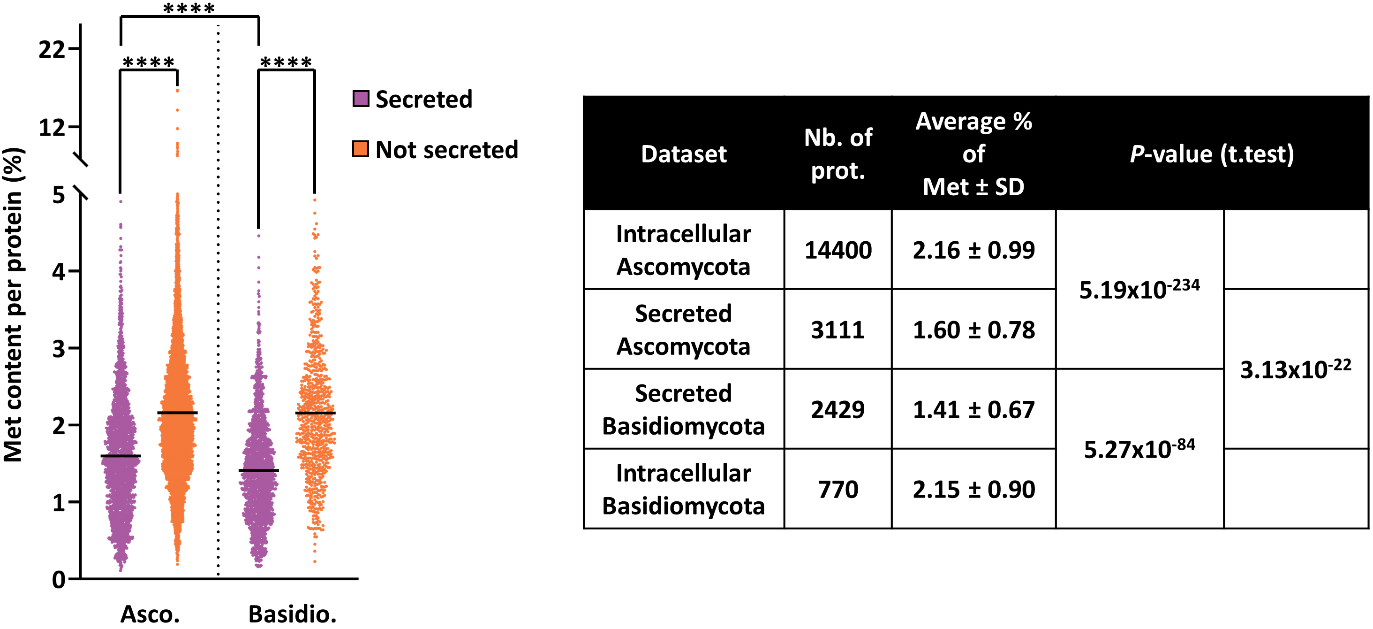

**Figure S2. Met content of Ascomycota and Basidiomycota secreted proteins and comparison with intracellular proteins**. Lists of proteins are in **Dataset 2**. Mean percentage of Met are indicated with *black* lines. Statistical analysis was performed by unpaired t-test with Welch’s correction (*****P* ≤ 0.001).

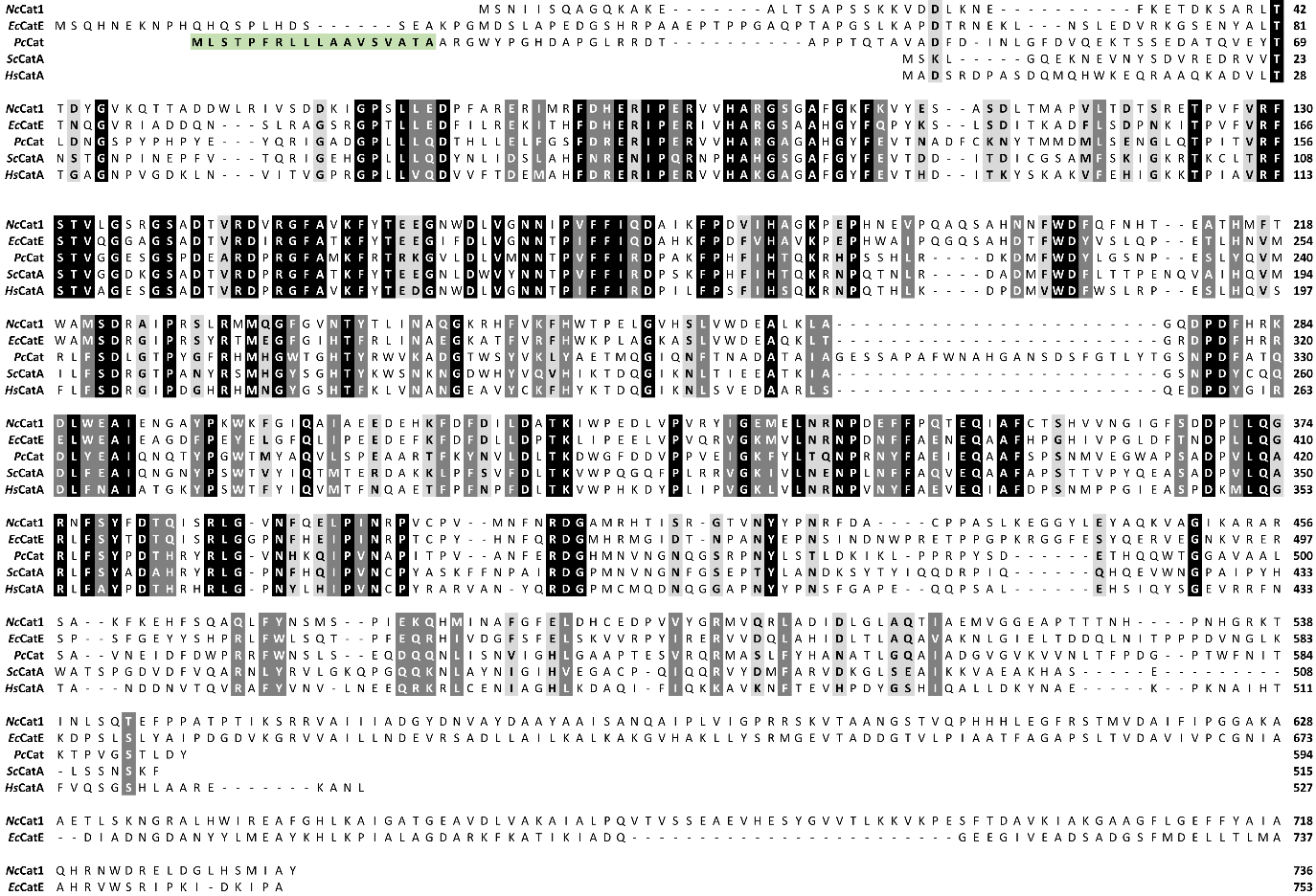

**A**

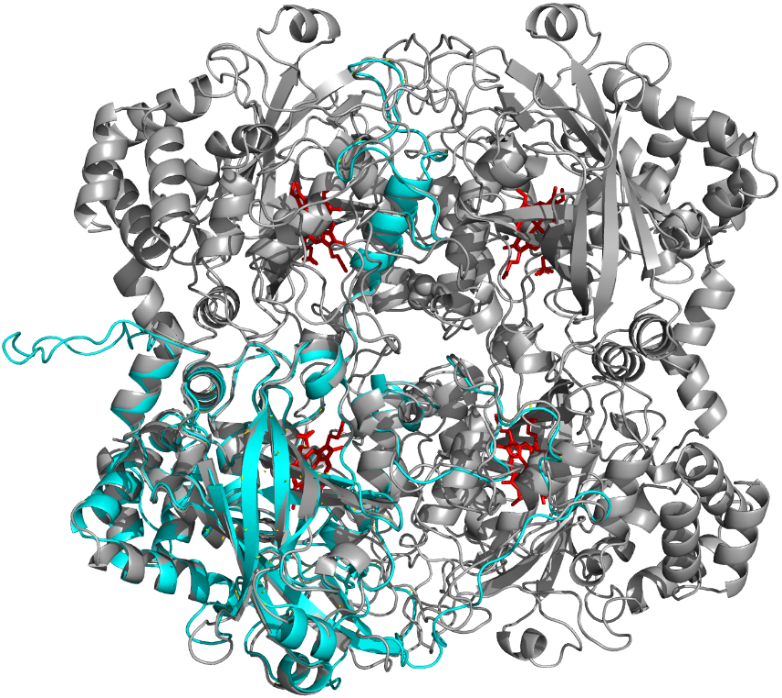

**B**

**Figure S3. Sequence and structure alignments of the *P. cinnabarinus* secreted catalase**. **A)** *P. cinnabarinus* secreted catalase protein sequence (*Pc*Cat; Uniprot# A0A060STN2) was aligned with *Neurospora crassa* Cat1 (*Nc*Cat1; Uniprot# Q9C168), *Escherichia coli* CatE (*Ec*CatE; Uniprot# P21179), *Saccharomyces cerevisiae* CatA (*Sc*CatA; Uniprot# P15202) and *Homo sapiens* CatA (HsCatA; Uniprot# P04040) using ClustalOmega (<https://www.ebi.ac.uk/Tools/msa/clustalo/>). *Black*, *dark* and *light grey* backgrounds represent strictly conserved, very similar and similar amino acids, respectively. *Green* background highlight secretion peptide signal. **B)** 3D model of the catalase A0A060STN2 (*cyan*) was downloaded from Alphafold database and aligned on the experimentally determined tetrameric structure of *S. cerevisiae* CatA (PDB #1A4E) (*grey*). Heme is represented in *red*. Structure manipulation and image generation were done using PyMol 2.3.2 (<https://pymol.org/2/>).

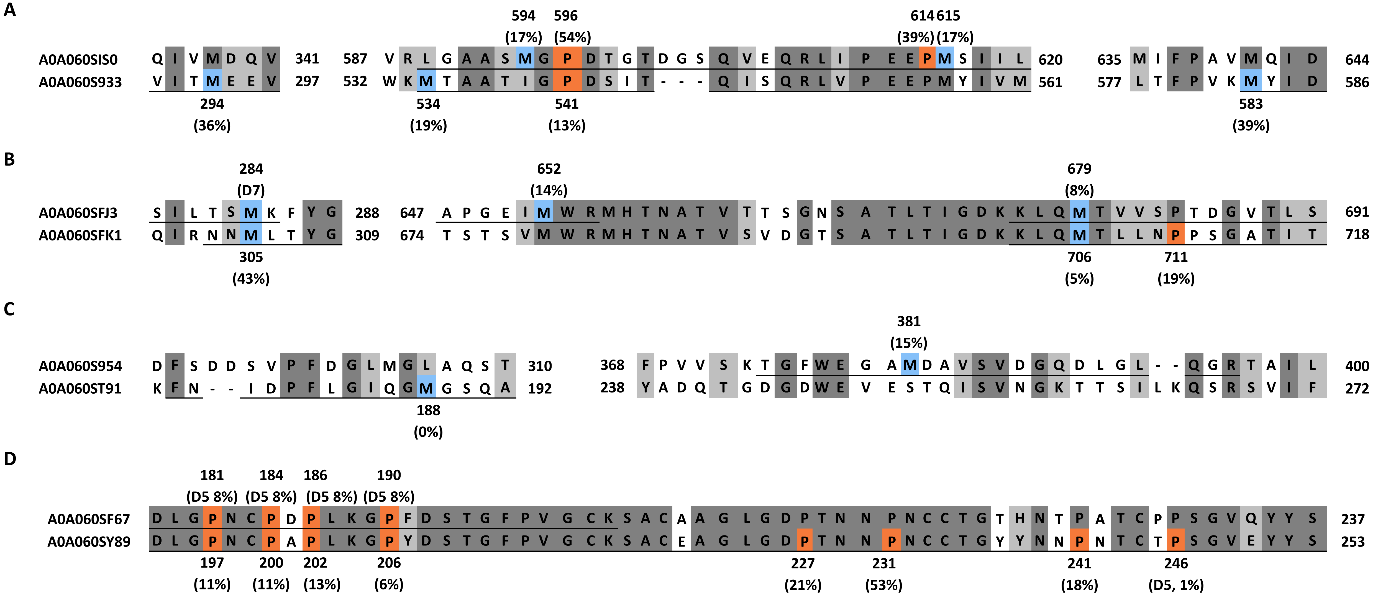

**Figure S4. Partial sequence alignment of GH16_2 (A), PL35 (B), protease (C) and GH152 (D) with identified oxidized residues.** Met and Pro found oxidized are on *blue* and *orange* background, respectively. *Dark* and *light grey* backgrounds represent strictly conserved and similar amino acids, respectively. Position and percentage of oxidation of the residues are indicated above or under the sequence. For Met, the percentage presented correspond to sum of Met^16^O and Met^16^O_2_ found at day 3, except indicated with ‘D7’, corresponding to Met found oxidized with ^18^O at day 7. For Pro, the percentage presented correspond to the sum of all oxidized forms, either ^16^O or ^18^O at day 3, excepted indicated with ‘D5’, corresponding to Pro oxidation found at day 5. Underlined sequences represent the peptides detected by LC-MS/MS. Accession numbers are on the left of the sequences. All data are presented in **Datasets 3** and **4**.

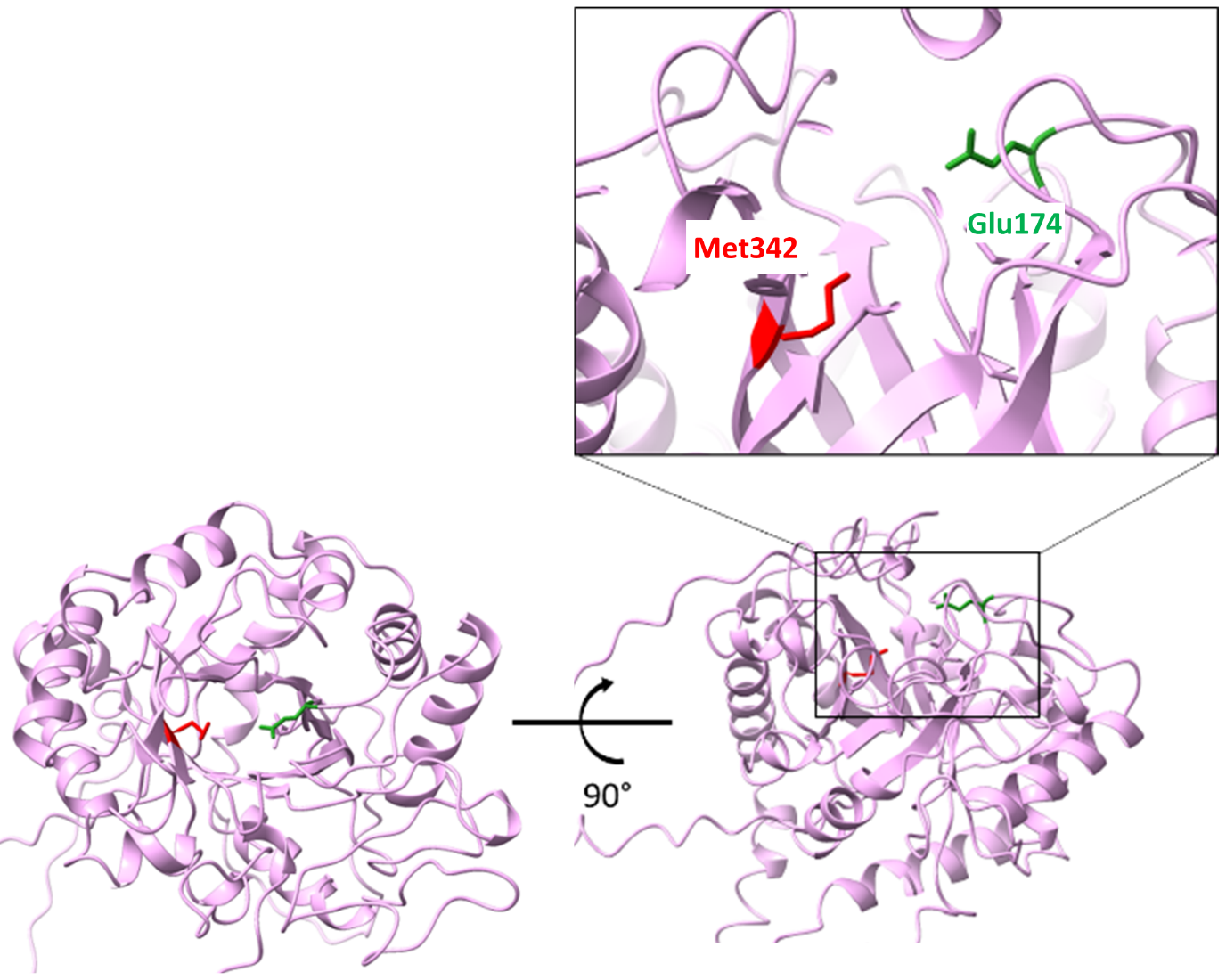

**Figure S5. 3D model of GH18 highlighting oxidized and conserved Met.** 3D model of the GH18 A0A060SC99 was downloaded from Alphafold database (<https://alphafold.ebi.ac.uk/entry/A0A060SC99>). The Met 342 found oxidized and the catalytic Glu 174 are represented as *red* and *green* sticks, respectively.

**Table S1. Met content in secreted and non-secreted proteins of *P. cinnabarinus*, fungi, metazoans, and plants.** The list of individual proteins is provided in the **Dataset 2**.

| Dataset | Description | Total nb. of organisms | Total nb. of prot. | Met percentage | | | | |
| --- | --- | --- | --- | --- | --- | --- | --- | --- |
|  |  |  |  | **Average** | ***P*-value (t.test)** | **Min** | **Max** | **Med.** |
| 2A | *P. cinnabarinus* predicted secreted proteins | 1 | 666 | 1.55 ± 0.82 | 2.30x10^-49^ | 0.24 | 5.88 | 1.43 |
| 2B | *P. cinnabarinus* predicted non secreted proteins | 1 | 9649 | 2.07 ± 0.96 |  | 0.18 | 23.16 | 1.97 |
| 2C | Fungal secreted proteins | 121 | 5542 | 1.52 ± 0.74 | 0^a^ | 0.11 | 7.41 | 1.44 |
| 2D | Fungal non secreted proteins | 400 | 15506 | 2.17 ± 0.99 |  | 0.18 | 16.67 | 2.04 |
| 2E | Metazoan secreted proteins | 1884 | 22164 | 2.68 ± 1.40 | 1.53x10^-132^ | 0.03 | 21.85 | 2.44 |
| 2F | Metazoan non secreted proteins | 738 | 30832 | 2.41 ± 1.07 |  | 0.07 | 18.03 | 2.27 |
| 2G | Plant secreted  proteins | 69 | 2960 | 2.59 ± 1.55 | 0.125 | 0.15 | 16.46 | 2.28 |
| 2H | Plant non secreted proteins | 1172 | 21013 | 2.55 ± 1.14 |  | 0.22 | 17.16 | 2.38 |

**^a^** The *P*-value being very close to 0, the calculation software cannot display the precise value.

**Table S2. Levels of Met oxidation in the secretomes obtained at day 3, 5 and 7.** The levels of non-oxidized and oxidized Met were calculated using all detected spectra (**Dataset 3B**).

|  | Day 3 | Day 5 | Day 7 |
| --- | --- | --- | --- |
| Number of identified proteins with detected Met | 75 ± 4 | 87 ± 3 | 86 ± 11 |
| Total spectral counts of detected Met | 749 ± 14 | 859 ± 73 | 913 ± 118 |
| *Spectral count* |  |  |  |
| Not modified Met | 136 ± 25 | 535 ± 34 | 542 ± 67 |
| Met^16^O | 122 ± 11 | 307 ± 40 | 351 ± 53 |
| Met^16^O_2_ | 15 ± 3 | 8 ± 3 | 12 ± 2 |
| Met^18^O | 459 ± 25 | 8 ± 2 | 5 ± 4 |
| Met^18^O_2_ | 17 ± 3 | 1 ± 1 | 2 ± 1 |
| *Percentage* |  |  |  |
| Not modified Met | 18.1 ± 3.0 | 62.3 ± 1.3 | 59.4 ± 0.4 |
| Met^16^O | 16.3 ± 1.5 | 35.6 ± 1.6 | 38.4 ± 0.9 |
| Met^16^O_2_ | 2.0 ± 0.3 | 1.0 ± 0.3 | 1.4 ± 0.1 |
| Met^18^O | 61.3 ± 3.9 | 0.9 ± 0.2 | 0.6 ± 0.5 |
| Met^18^O_2_ | 2.3 ± 0.4 | 0.2 ± 0.1 | 0.3 ± 0.1 |

*SC*, spectral counts; Met^16^O, methionine sulfoxide with ^16^O; Met^16^O_2_, methionine sulfone with ^16^O; Met^18^O, methionine sulfoxide with ^18^O; Met^18^O_2_, methionine sulfone with ^18^O.

**Table S3. Levels of Pro oxidation in the secretomes obtained at day 3, 5 and 7.** The levels of non-oxidized and oxidized Pro were calculated using all detected spectra (**Dataset 4B**).

|  | Day 3 | Day 5 | Day 7 |
| --- | --- | --- | --- |
| Number of identified proteins with detected Pro | 130 ± 3 | 147 ± 3 | 143 ± 15 |
| Total spectral counts of Pro | 3099 ± 145 | 3150 ± 197 | 2899 ± 196 |
| *Spectral count* |  |  |  |
| Not modified Pro | 2866 ± 146 | 3017 ± 193 | 2794 ± 195 |
| Glutamic 5-semialdehyde-^16^O (SC) | 75 ± 9 | 61 ± 3 | 57 ± 3 |
| Pyroglutamic acid-^16^O (SC) | 47 ± 7 | 50 ± 3 | 35 ± 7 |
| Sum-^16^O (SC) | **122 ± 5** | **112 ± 1** | **92 ± 10** |
| Glutamic 5-semialdehyde-^18^O (SC) | 104 ± 8 | 16 ± 4 | 11 ± 4 |
| Pyroglutamic acid-^18^O (SC) | 7 ± 4 | 5 ± 2 | 3 ± 1 |
| Sum-^18^O (SC) | **111 ± 6** | **20 ± 5** | **14 ± 4** |
| *Percentage* |  |  |  |
| Not modified Pro | 92.4 ± 0.4 | 95.8 ± 0.2 | 96.4 ± 0.4 |
| Glutamic 5-semialdehyde-^16^O (%) | 2.4 ± 0.3 | 2.0 ± 0.2 | 2.0 ± 0.1 |
| Pyroglutamic acid-^16^O (%) | 1.5 ± 0.2 | 1.6 ± 0.1 | 1.2 ± 0.2 |
| Sum-^16^O (%) | **3.9 ± 0.1** | **3.6 ± 0.2** | **3.2 ± 0.3** |
| Glutamic 5-semialdehyde-^18^O (%) | 3.4 ± 0.3 | 0.5 ± 0.1 | 0.4 ± 0.1 |
| Pyroglutamic acid-^18^O (%) | 0.2 ± 0.1 | 0.1 ± 0.1 | 0.1 ± 0.1 |
| Sum-^18^O (%) | **3.6 ± 0.3** | **0.6 ± 0.1** | **0.5 ± 0.2** |

**Table S4. Abundance of detected H_2_O_2_-consuming enzymes in the secretomes obtained at day 3, 5 and 7.** The %NSAF were calculated using all detected spectra for each protein (**Dataset 1**).

| Uniprot Accession | %NSAF ± SD | | |
| --- | --- | --- | --- |
|  | **Day 3** | **Day 5** | **Day 7** |
| *Catalase* |  |  |  |
| A0A060STN2 | 0.02 ± 0.02 | 0.33 ± 0.05 | 0.42 ± 0.01 |
| *Class II peroxidase* |  |  |  |
| A0A060S827 | *n.d.* | 0.01 ± 0.02**^a^** | 0.03 ± 0.06**^a^** |
| A0A060SDU0 | *n.d.* | 0.01 ± 0.01**^a^** | *n.d.* |
| A0A060SJB5 | *n.d.* | *n.d.* | 0.07 ± 0.12**^a^** |
| A0A060SRR4 | *n.d.* | *n.d.* | 0.03 ± 0.05**^a^** |
| A0A060SYJ1 | 0.12 ± 0.01 | 0.18 ± 0.00 | 0.09 ± 0.02 |
| *AA9-LPMO* |  |  |  |
| A0A060S8F5 | *n.d.* | *n.d.* | 0.02 ± 0.02 |
| A0A060S9V0 | 0.02 ± 0.02**^a^** | 0.01 ± 0.01**^a^** | 0.03 ± 0.05**^a^** |
| A0A060SPU4 | *n.d.* | *n.d.* | 0.01 ± 0.02**^a^** |
| A0A060SQW9 | *n.d.* | *n.d.* | 0.01 ± 0.02**^a^** |

**^a^** High SD value is because the protein was not detected in all triplicates. *n.d., not detected.*

**Table S5. Number of predicted secreted catalases in fungi.** The protein sequence of *P. cinnabarinus* secreted catalase (A0A060STN2) was used to search for homologs by BLASTP against all fungal genomes. Among the 9492 hits, 1630 were predicted to be secreted.

| Phyla | Nb. Genome | Nb. Secreted Catalase | Secreted catalase per genome |
| --- | --- | --- | --- |
| Ascomycota | 1069 | 1182 | 1.11 |
| Basidiomycota | 256 | 374 | 1.46 |
| *White rot* | *85* | *101* | *1.19* |
| *Ectomycorrhizal* | *59* | *73* | *1.24* |
| *Other saprotroph* | *40* | *69* | *1.73* |
| *Phytopathogen* | *34* | *85* | *2.50* |
| *Brown rot* | *24* | *29* | *1.21* |
| *Other* | *14* | *17* | *1.21* |
| Mucoromycota | 48 | 70 | 1.46 |
| Chytridiomycota | 3 | 3 | 1.00 |
| Zoopagomycota | 1 | 1 | 1.00 |
| Cryptomycota | 0 | 0 | 0.00 |
| Total | 1377 | 1630 | 1.20 ± 0.24 |
